# Amorphous calcium phosphate-coated surfaces as a model for bone microenvironment in prostate cancer

**DOI:** 10.1101/2023.03.20.533462

**Authors:** Rebeca San Martin, Priyojit Das, Tianchun Xue, Morgan Rose Brown, Renata Dos Reis Marques, Michael Essington, Adrian Gonzalez, Rachel Patton McCord

## Abstract

Bone metastasis remains one of the biggest challenges in the treatment of prostate cancer, and other solid tumors such as breast, lung, and colon. Modeling a complex microenvironment in-vitro, such as the bone niche, requires interrogation of cell-cell interactions, specific extracellular matrix proteins and a high calcium environment. Here, we present a fast and cost-effective system in which commercially available, non-adhesive, cell culture vessels are coated with amorphous calcium phosphate (ACP) as a surrogate for bone matrix. We further present modified protocols for subculturing cells, as well as nucleic acid and protein collection in high calcium samples. We find that prostate epithelial cell lines show increased adhesion and proliferation when cultured in these surfaces, as well as independence from androgen starvation. We observe gene expression changes on ACP surfaces in early adenocarcinoma cell lines which may reflect alterations relevant to prostate cancer progression.

**Summary statement:** To model the role of calcium in the microenvironment of the metastatic bone niche, we developed a cost-effective way to coat cell culture vessels in bioavailable calcium, and show that it has an effect on prostate cancer cell survival

## Introduction

Prostate cancer remains a challenge in healthcare in the United States since it is the most commonly diagnosed malignancy in men (Siegel et al., 2022). Patients diagnosed with localized prostate cancer can present with circulating tumor cells (Broncy and Paterlini-Brechot, 2019; Theil et al., 2021; Todenhofer et al., 2016). Once these cells escape the primary tumor, they can potentially invade other tissues and develop into micro metastases. In particular, prostate cancer metastasizes preferentially to bone through organotropic mechanisms that are poorly understood. In the bone microenvironment, these metastatic foci associate with the hematopoietic stem cell niche (Shiozawa et al., 2008; Shiozawa et al., 2011) and can remain dormant for extended periods. In this context, the endosteal layer of quiescent osteoblasts induces the dormant phenotype via several mechanisms (Taichman et al., 2013; Tanaka et al., 2021). Importantly, this bone microenvironment is the primary repository of calcium, which is incorporated and mobilized from the osteoid matrix to respond to systemic demand (reviewed in Murshed, 2018; Tresguerres et al., 2020). Factors like race (Noel et al., 2021), age (I.O.F., 2022; Veldurthy et al., 2016), and concomitant morbidities like diabetes (Ahn et al., 2017; Cipriani et al., 2020; Jackson and Moseley, 2020; Teissier et al., 2022) impact bone calcium homeostasis: they can result in osteopenia (loss of calcified bone mineral density) and osteoporosis (loss of mineral density that results in decreased bone strength). Adding complexity to these phenotypes, the standard of care for prostate cancer also negatively impacts bone matrix integrity. Specifically, androgen deprivation therapy (ADT: chemical castration, testosterone blockade) interferes with osteoblast homeostasis resulting in loss of mineral density (Badri et al., 2019; Tsaur et al., 2018). These events could create a perfect storm of a microenvironment rich in bioavailable calcium. In this study, we set out to model the behavior of prostate-derived cancer cells in such an environment.

Unfortunately, the COVID-19 pandemic placed an unusual strain on the supply chain of polypropylene-derived plastics necessary for research (Press, 2021; Sheridan, 2021). As a result, the production of cell culture specialty vessels like those needed to model the bone microenvironment was severely impacted. Spurred by this uncertainty in sourcing material for our studies, we set out to modify an amorphous calcium phosphate (ACP) chemical deposition system, originally developed to coat titanium for use on bioimplants (Kim et al., 2013). Our system is useful for depositing amorphous calcium phosphate on untreated plastic cell culture vessels and glass coverslips. The technique is cost-effective, modular, and robust and can generate hundreds of cell culture plates in a matter of weeks, ensuring continuity of research independent of supply issues. Here we describe the advantages and challenges of the system, along with the methods developed for culturing prostate cancer cell lines on these surfaces and deriving viable biological samples from them.

Culturing normal (RWPE), adenocarcinoma (LNCaP), and bone metastatic (VCaP, MDAPCa2a) prostate epithelium on these ACP-coated vessels, we characterized an increased adherent affinity to calcium-rich surfaces and an induction of proliferation. We further identified changes in the transcription of genes related to cell proliferation and metastasis progression. Intriguingly, the presence of calcium compensated for the absence of testosterone in some cases, providing new insight into an alternative mechanism for the rise of castration resistance.

## Results

### Alkali-treated titanium serves as a catalyst for amorphous calcium phosphate aggregate formation and a surface for coating

To develop a cell culture system in which prostate cancer cells interact with a high bioavailable calcium microenvironment, similar to the metastatic bone site, we implemented a modified version of the amorphous calcium phosphate (ACP) coating on alkali-treated titanium as previously described (Kim et al., 2013). We scaled the protocol to activate and coat multiple titanium plates of larger sizes, amenable for cell culture in 6 well plates (Figure 1A). Activating the titanium surfaces in five molar potassium hydroxide results in mild etching of the plate surface, evident under scanning electron microscopy (SEM) (Fig. S1 A-B). Incubation of these activated surfaces in ACP solution results in aggregate formation, visible on the supernatant of the solution, as early as 24 hours after the start of the coating phase (Figure 1B). Replacing two-thirds of the solution every 24 hours allowed for less disturbance of the coating surface and faster nucleation of the new solution, resulting in a white, chalky coating after four days of incubation (Figure 1C), which persists after washing with chemically pure water. An evaluation of the coating using SEM confirms an amorphous, filamentous structure similar to that previously reported (Figure 1D) (Kim et al., 2020; Vaquette et al., 2013)

**Figure 1.**
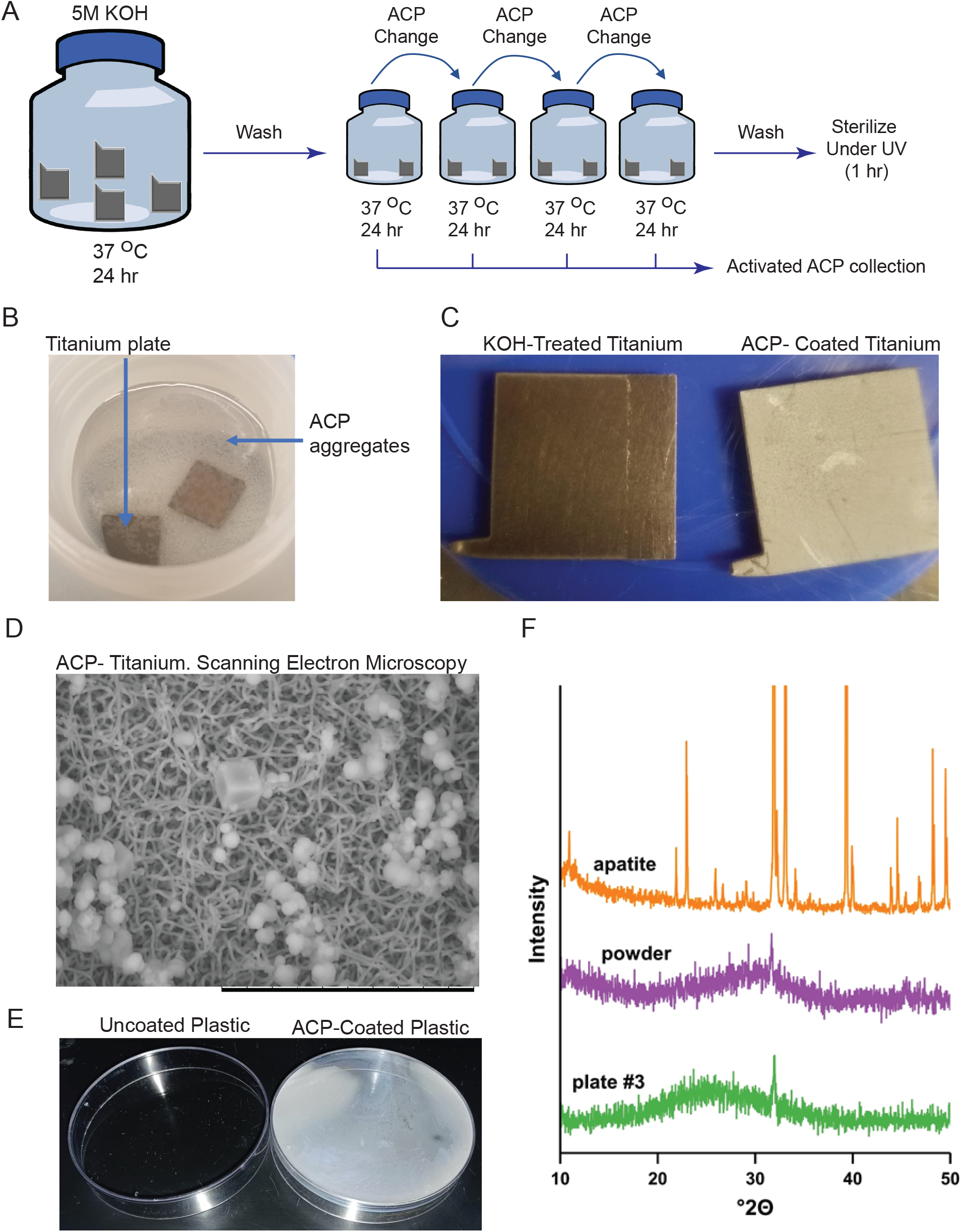
Coating of potassium hydroxide-activated titanium plates with amorphous calcium phosphate (ACP) A. Graphic representation of the method for amorphous calcium deposition (ACP) on titanium surfaces and collection of activated ACP solution for plastic coating B. Incubation of potassium hydroxide-activated titanium plates with ACP solution for 24 hours is enough to elicit nucleation of precipitate in the solution. C. Potassium hydroxide-activated titanium plates conserve a metallic sheen (left), while plates exposed to ACP solution for four days are coated with a persistent, white, chalky substance (right) D. Scanning electron microscopy shows that ACP-coated titanium plates are coated with a combination of amorphous globular structures and a fibrillar-like network (scale: 20 microns). Sodium chloride crystals are also observed, interspersed among those structures (cuboidal structure). E. ACP coating is persistent on non-treated plastic surfaces. ACP-coated plate (right) shows characteristic white powdered deposition compared to control plates incubated in buffer (left). F. X-ray diffraction (XDR) analysis reveals two peaks in the powdered ACP collected from petri dishes (purple-“powder”): a weak peak at 31.67 °2θ (d = 2.79 Å) and a weaker peak at 45.5 °2θ (d = 1.99 Å), while the sample coated on glass coverslips (green-“plate #3”) shows profiles of a single weak diffraction line (peak) at 31.90 °2θ (d = 2.81 Å). The graph compares the XRD profiles of the unknown precipitates to that of apatite.

### X-ray diffraction analysis of ACP aggregates

The material deposited onto three ACP-coated glass coverslips, and 500 mg of ACP precipitate, collected from eighteen Petri dishes that were coated and then scraped as described in the methods section, were evaluated via X-ray diffraction. The three glass-coated coverslips generated similar XRD profiles (Figure. 1E – green); these profiles consisted of a single weak diffraction line (peak) at 31.90 °2θ (d = 2.81 Å). The powder sample (Figure 1E – Purple) generated two peaks: a weak peak at 31.67 °2θ (d = 2.79 Å) and a weaker peak at 45.5 °2θ (d = 1.99 Å). Figure 1F compares the XRD profiles of the unknown precipitate to that of hydroxyapatite. Based on the XRD analysis, the only crystalline substances in the samples are KCl and NaCl. The bulk of the precipitate appears to consist of an amorphous (microcrystalline) substance, similar to amorphous calcium phosphate reported in the literature.

### Activated ACP solution can be used to coat plastic and glass cell culture vessels

To evaluate how prostate cancer cells interact with calcium-coated surfaces, non-cell culture-treated plastic vessels were coated with activated ACP solution immediately after the four-day pooling from the titanium coating phase. We used bacterial culture Petri dishes, non-treated six-well plates, or glass coverslips to prevent confounding adhesion interactions with treated plastics. Treatment with ACP for four days at 37°C is enough to coat both types of plastic (Figure 1E) or glass (Fig. S1C). This coating is persistent after washing in chemically pure water, with vigorous shaking. Based on the collection of ACP precipitate used for the X-ray diffraction studies, and given that the coating appears to be homogeneous, we estimate that each ten-centimeter plate is coated with about twenty-seven milligrams of ACP or about 0.4 mg/cm^2^.

As described in the methods, sterilization under ultraviolet light does not affect the ACP structure or coating and is sufficient to prevent contamination during cell culture.

Petri dishes coated in ACP were incubated in standard full media formulations used for prostate cancer cell lines (Table S1) to verify the stability of the calcium phosphate coating during cell culture. We find that chemically defined media that does not contain serum (KSF) can strip a significant amount of ACP from the surfaces. To further evaluate changes in calcium concentration in the media, media pooled from four-day incubation on ACP-coated surfaces were submitted to ion chromatography analysis at the Water Quality Core Facility of the University of Tennessee, Knoxville. These results (Fig. S1D) confirm the visual observation of ACP stripping in KSF. Some increase in calcium concentration occurred on HPC1 media, with the coating still visible under the microscope.

### Culturing prostate cancer cell lines on ACP-coated surfaces fosters their adhesion, proliferation, and independence from androgen depletion

We cultured the adenocarcinoma cell line LNCaP and bone metastatic cell lines VCaP (White-European) and MDAPCa2a (African American) on ACP-coated titanium plates and plastics to evaluate the influence of an environment rich in bio-available calcium on prostate cancer epithelium.

The adenocarcinoma cell line (LNCaP) attaches readily to the ACP-coated surfaces (Figure 2A). The high attachment affinity for the coating requires a sequence of EDTA chelation, extended trypsin incubation, and aggressive shaking for subculturing, as described in the Methods. Cell viability is not affected by culturing on ACP surfaces, but the presence of both ACP and testosterone (DHT) supplementation increases proliferation (Figure 2A, right). Strikingly, LNCaP cells cultured in ACP-coated vessels are not sensitive to the absence of testosterone modeled using charcoal-dextran-treated serum in the culture media. After four days of culture on normal plastic and charcoal-dextran serum conditions, we observe a well-documented decrease of proliferation in LNCaP while cells proliferate at the normal rate in ACP conditions.

**Figure 2.**
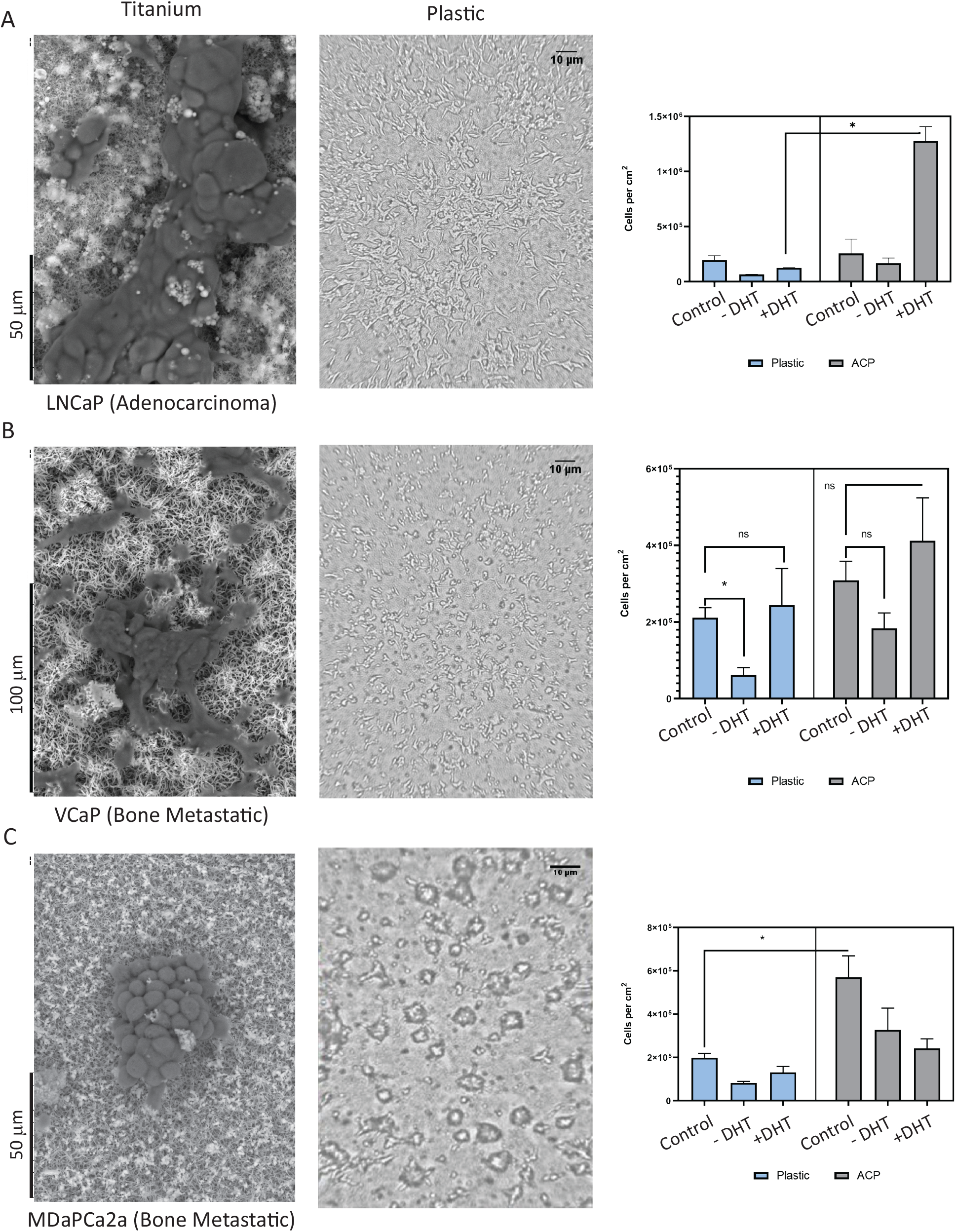
Epithelial cell culture on ACP-coated, non-adhesive plastic surfaces. A. Prostate adenocarcinoma cell line LNCaP grown on amorphous calcium phosphate surfaces. Scanning electron microscopy of cells grown in ACP-titanium coated surfaces, bar 50 μm (left). Cells cultured on ACP coated-non adhesive plastic for 48 hours (middle). Cell culture yield from full media control, testosterone depleted (-DHT), or testosterone supplemented (+DHT) conditions. B. Bone metastatic cell line VCaP grown in amorphous calcium phosphate surfaces. Scanning electron microscopy of cells grown in ACP-titanium coated surfaces, bar 100 μm (left). Cells cultured on ACP coated-non adhesive plastic for 48 hours (middle). Cell culture yield from full media control, testosterone depleted, or testosterone supplemented conditions. C. Bone metastatic cell line MDAPCa2a grown in amorphous calcium phosphate surfaces. Scanning electron microscopy of cells grown in ACP-titanium coated surfaces, bar 50 μm (left). Cells cultured on ACP coated-non adhesive plastic for 48 hours (middle). Cell culture yield from full media control, testosterone depleted, or testosterone supplemented conditions.

The bone metastatic cell VCaP of European ancestry adheres to ACP surfaces with a high affinity that requires modified subculturing protocols as described. (Figure 2B). VCaP cells show a prototypical response to the absence of testosterone in normal plastic at four days, with a significant proliferation decrease when cultured in media containing charcoal dextran-treated serum. This effect is rescued by the addition of testosterone to the media. Similarly to our observations in LNCaP, culturing the VCaP cell line on ACP-coated surfaces bypasses the proliferation arrest that results from the absence of testosterone (Figure 2B right).

In contrast to the other cells used in this study, the bone metastatic cell MDAPCa2a cannot adhere to cell culture-treated plastics unless coated with a proprietary mixture of fibronectin and collagen (FNC coating - Athena biologicals). Growing on this extra-cellular matrix protein context and at the 4-day experimental timepoint, MDAPCa2a cells show a small proliferative decrease when cultured in media that contains a charcoal dextran-treated serum that, similarly to what occurs in VCaP, is rescued by the addition of testosterone. Strikingly, MDAPCa2a culture on ACP-coated surfaces (titanium and plastic) does not require FNC coating. Cells adhere to the surface, making both the characteristic 3D colonies observed in plastic, with single cells growing in a monolayer as well (Figure 2C). Further, these cells show a similar trend as VCaP regarding the presence and absence of testosterone in plastic (Figure 2C right). The presence of the ACP surface results in a significant increase in the proliferation of MDAPCa2a cells in the presence of complete media. In contrast to VCaP cells, however, MDAPCa2a cells show a non-significant decrease in proliferation when cultured in media containing charcoal-dextran-treated serum. Interestingly, this effect is not rescued by the addition of testosterone, suggesting that the absence of another steroid-like compound present in the serum could affect the proliferation of these cells (Figure 2C right).

### Culture in ACP-coated surfaces results in an altered transcriptional landscape

To evaluate the transcriptional effect that bioavailable calcium coating has on prostate epithelial cells, we collected RNA from our model cell lines (RWPE, LNCaP, VCaP, and MDAPCa2a) cultured under different conditions: cell culture-treated plastic or ACP-coated non-cell culture treated vessels were used in combination with control media, media without testosterone (using charcoal-dextran stripped serum), or media supplemented with testosterone (in charcoal-dextran stripped serum). We found that RNA quantity and quality remained high when isolated from cells grown on ACP surfaces (Fig. S1E) and we were able to construct and sequence RNA-seq libraries from all samples.

Two unsupervised analyses were used to compare the transcriptomic output of each of these experimental conditions. First, genes that showed a log2 fold change with a cutoff of +/- 1 for each condition were analyzed using Metascape (Zhou et al., 2019) (Full gene lists and enrichment results available in Data S1). Gene ontology categories derived from these outputs were manually clustered for similarities, and common gene lists were compiled. In agreement with our proliferation data, this analysis revealed that three main gene ontology terms are enriched for transcriptional upregulation when non-tumorigenic (RWPE) or early adenocarcinoma (LNCaP) cell models are grown in contact with calcium surfaces: Cell cycle progression (Figure 3A, Table S2), oxidative phosphorylation (Figure 3B, Table S2) and translation/ribosome assembly (Fig. S2). Genes common to these ontologies and comparisons include CDK1, COMMD3, B2M, CDK7, and COX17.

**Figure 3.**
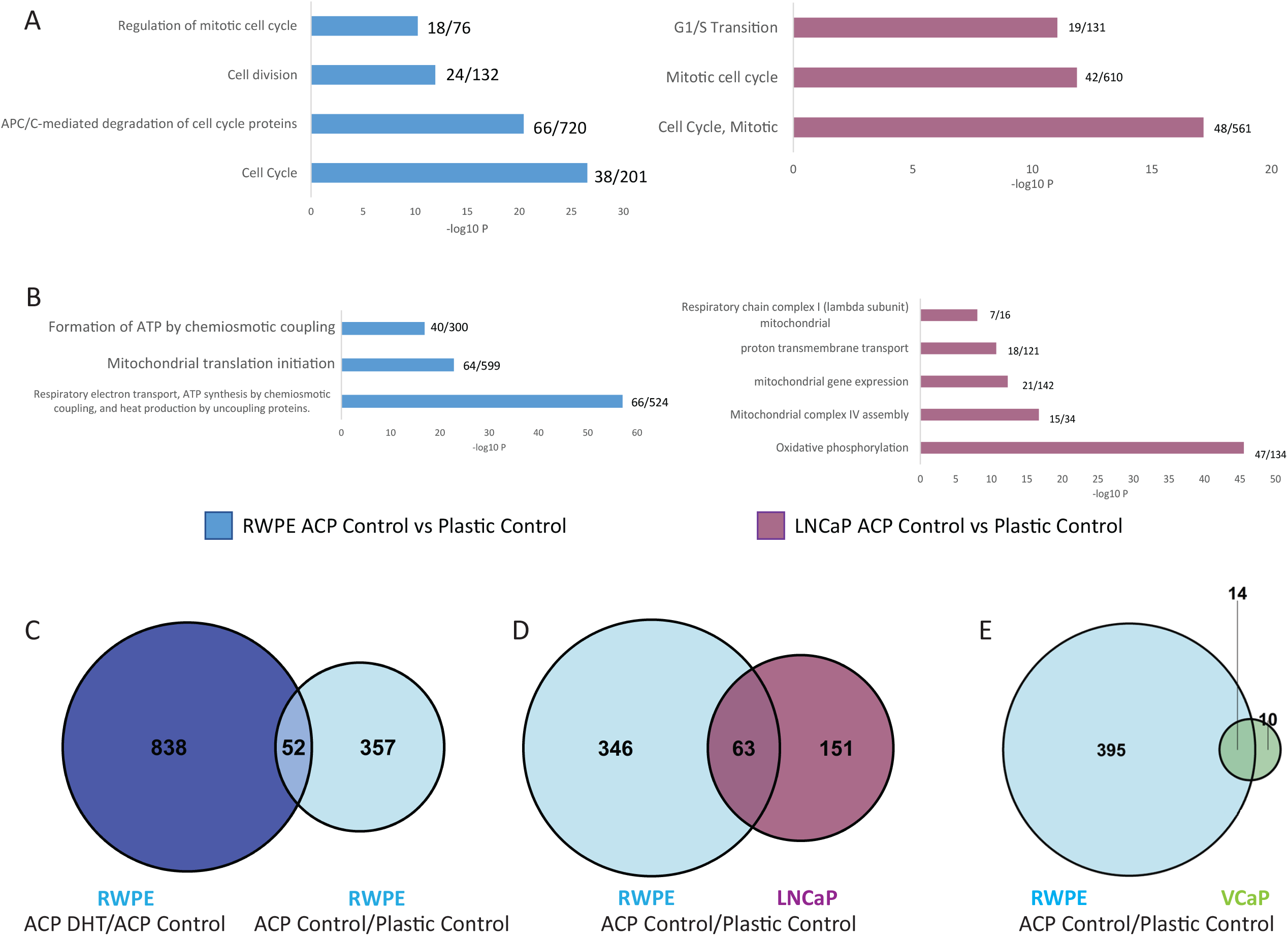
Congruent changes in transcription are observed in prostate non-tumorigenic epithelium and adenocarcinoma cells cultured on ACP surfaces. A. Culturing the non-tumorigenic cell line RWPE (left) and the early adenocarcinoma cell line LNCaP (right) on calcium-coated surfaces induces transcription of genes related to cell cycle progression when compared to culture in normal plastic. (Full gene list in Table S2) B. Culturing RWPE and LNCaP on calcium-coated surfaces induces transcription of genes related to oxidative phosphorylation. (Full gene list in Table S2) C. The addition of testosterone to RWPE culture on calcium-coated surfaces further enhances transcription of a cohort of 52 genes. D. Comparison of transcriptionally upregulated genes common between normal epithelium (RWPE), adenocarcinoma (LNCaP) and (E) bone metastatic cell (VCaP) line, shows common hubs of genes that are upregulated in calcium-rich culture conditions (Gene lists: Table S3)

Further, to assess whether common genes were affected in different cell lines treated in the same conditions, we intersected gene lists that showed a significant change (FDR p-adjusted value <0.001). We find that fifty-two genes upregulated by culturing RWPE on calcium-rich surfaces are further upregulated when culture on calcium is supplemented with testosterone (Figure 3C). Sixty-three genes that are upregulated in normal epithelium cultured on calcium are also upregulated in adenocarcinoma cell lines under the same condition (Figure 3D). Genes common to these comparisons include CDK1, PARPBP, ANKRD22, ITGAE, and COX7B. Only fourteen genes are common among the normal epithelium-bone metastatic cell line comparison (Figure 3E, Table S3).

Our recent work has shown that there are concerted changes in the 3D genome architecture related to compartment identity that could be predictive of malignancy status (San Martin et al., 2022). In this previous study, we identified a cohort of genes that appear in the spatial B compartment (heterochromatin) in normal epithelial cell lines that switch conformation to the A compartment (euchromatin) in cancer cell lines. This switch in compartment identity is sometimes accompanied by transcriptional upregulation. Using this list of genes as a reference, we asked whether transcriptional induction of these genes occurred due to contact with calcium surfaces (Fig. S3). We find that transcription of CDK1, DEPDC1B, and PARPBP is upregulated in normal epithelial cells as a response to calcium and testosterone (Figure 4A, yellow box).

**Figure 4.**
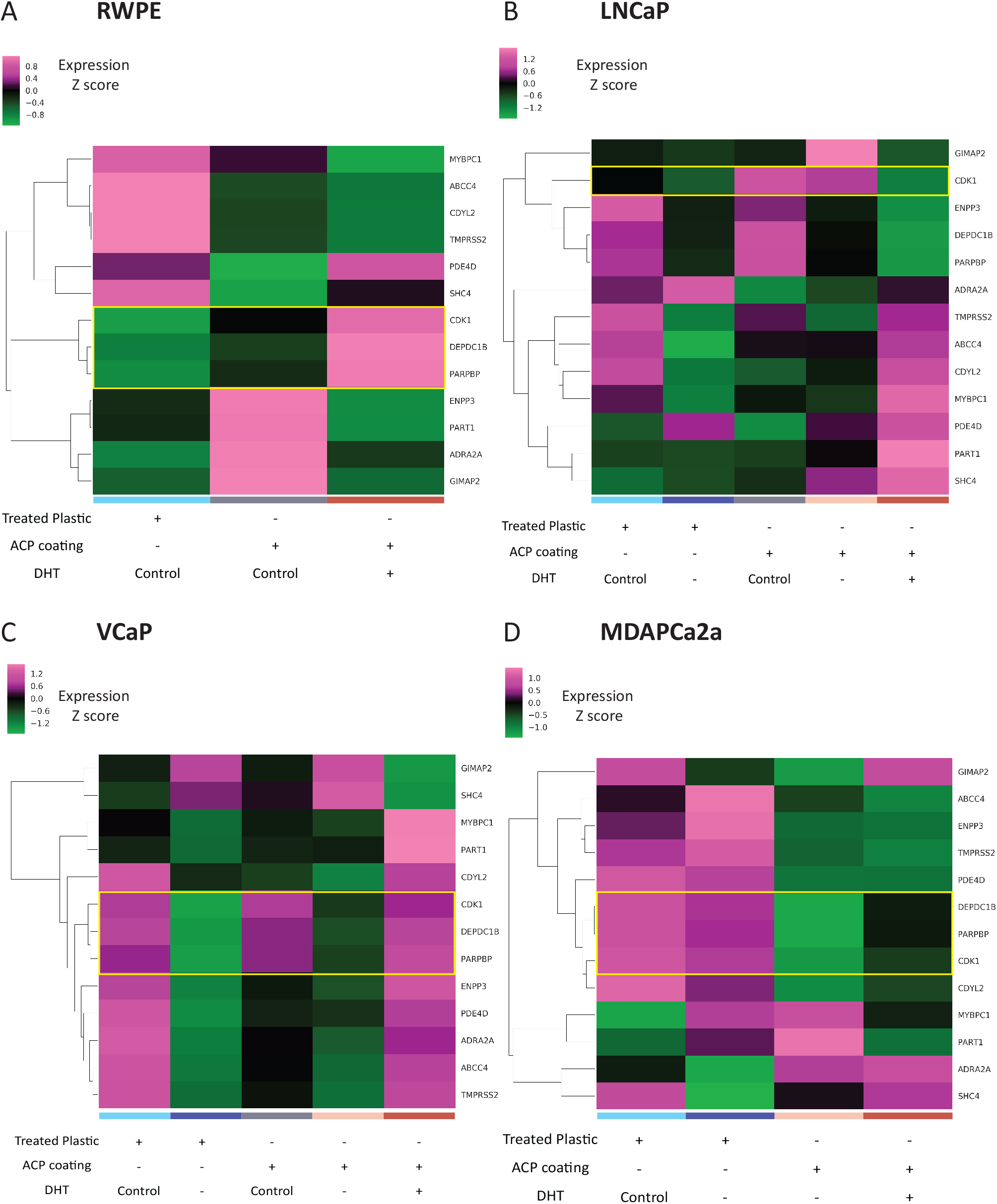
Transcription of genes that undergo 3D genome architecture changes in prostate cancer bone metastasis is affected by culturing in ACP-coated plastic vessels. A. Culturing the non-tumorigenic cell line RWPE on calcium-coated surfaces induces transcription of the CDK1, DEPDC1B and PARPBP genes in the presence of 100 mM testosterone supplementation (yellow rectangle). B. Culturing the adenocarcinoma cell line LNCaP on calcium-coated surfaces induces transcription of select genes. Among these genes, CDK1 transcription is upregulated (yellow rectangle) C. Transcription of the CDK1, DEPDC1B and PARPBP genes by the bone metastatic cell line VCaP is dependent on the presence of testosterone, and mildly induced by culturing on calcium coated surfaces (yellow rectangle). D. Culturing the bone metastatic cell line MDaPCa2a on calcium-coated surfaces represses transcription of CDK1, DEPDC1B and PARPBP, in contrast to the effect observed VCaP. This effect is not rescued by the addition of testosterone. All heat maps of RNA-seq analysis show per-gene z-score of batch effect corrected data, hierarchically clustered based on Euclidean distance and average linkage.

In contrast, MYBPC1, ABCC4, CDYL2, and TMPRSS2 transcription is downregulated by external calcium, regardless of testosterone supplementation (Figure 4A). The transcriptional induction of CDK1 is conserved in LNCaP cells cultured in calcium with both complete media and androgen depletion, but interestingly transcription is depressed when cells are supplemented with testosterone in calcium (Figure 4B). Contrary to RWPE, transcription of MYBPC1, ABCC4, CDYL2, and TMPRSS2 is upregulated in LNCaP cultured on calcium with testosterone supplementation (Figure 4B). In stark contrast, expression of CDK1, DEPDC1B, and PARPBP in the bone metastatic cell line VCaP is dependent on the presence of testosterone rather than calcium, while the presence of calcium inhibits their transcription in the bone metastatic cell line MDAPCa2a, regardless of the presence of testosterone (Figure 4C-D). Finally, transcription of MYBPC1, ABCC4, CDYL2, and TMPRSS2 is repressed by testosterone depletion in both plastic and calcium surfaces and calcium surfaces with control media but rescued by testosterone supplementation (100mM) in calcium culture. These genes are downregulated in MDAPCa2a in all calcium surface conditions. (Figure 4C-D).

Finally, we evaluated whether transcription of genes whose expression increased in the context of our experimental bioavailable calcium model also increase in patients by analyzing available prostate data from The Cancer Genome Atlas (GDC TCGA PRAD). This cohort includes data from six hundred and twenty-three patients (81% tumor samples, 19% normal samples). Using data from patients with a reported Gleason score, we find that CDK1 and DEPDC1B high expression is upregulated in tumor samples (Fig. S4A) and correlates with biochemical recurrence (Fig. S4B). Expression of these genes also correlates with a higher Gleason Score and is concurrent with each other as Gleason Score increases (FigS4C-D).

## Discussion

Advanced prostate cancer metastasizes preferentially to the bone (Bubendorf et al., 2000). Importantly, at the time of diagnosis, about a quarter of all patients already present with tumor-circulating cells (reviewed by Schilling et al., 2012; Theil et al., 2021). If these disseminated cancer cells were to arrive at the bone niche through circulation, they could remain in a dormant state under the influence of osteoblast-secreted signals (Shiozawa et al., 2010; Taichman et al., 2013).

Previous studies have characterized a reactive, fracture repair-like endosteum associated with trabecular prostate cancer metastasis (San Martin et al., 2017) and the important role of bone calcium in the context of cancer (Wang et al., 2018). These two separate observations can be intimately related since the processes of bone remodeling, via activation of both osteoblasts and osteoclasts, plays an important role in the release of bioavailable calcium (reviewed by Croucher et al., 2016)

The present work established a cell culture system incorporating a solid-phase bioavailable calcium coating to model a receptive bone microenvironment. In this context, bioavailable calcium is defined as A) calcium that is compatible with body fluids and is present in native tissues in solid form, B) fosters biological activity bioactivity by mediating adhesion and proliferation, and C) a matrix that can be degraded by cells and release calcium ions (Jeong et al., 2019). Specifically, amorphous calcium phosphate (ACP) coatings are biologically relevant since they foster new bone formation by osteoblasts, inducing faster mineralization and osteogenic induction (Vaquette et al., 2013). ACP also enhances MSC-mediated reconstruction of bone defects and engraftment (Brennan et al., 2021), and increases titanium implant biocompatibility and bonding strength, even in the presence of coating resorption (Maxian et al., 1993).

Further, it has been proposed that ACP acts as a potential intermediate in the precipitation of hydroxyapatite *in vitro* (Eanes et al., 1965; Edén, 2021). While there is controversy about the presence of non-crystalline phase apatite and amorphous calcium phosphate in bone (Boskey, 1997), several studies have characterized disordered calcium phosphate deposition. Amorphous calcium compounds are predominant in new segments of ray bones in zebrafish (Mahamid et al., 2008), where they act as precursors of the mature bone’s carbonated hydroxyapatite. Further, a detailed transformation mechanism of amorphous calcium phosphate particles into apatite-like crystals has also been described within the confined domains of a fibrous nanocomposite resembling the collagen matrix in the developing bone (Lotsari et al., 2018). Conflicting studies that characterized the absence of amorphous calcium phosphate content in bone (Kaflak-Hachulska et al., 2003) were hindered by a limited sample size of two adult cadaveric donors since ACP deposition is more prevalent in young vertebrates (Termine and Posner, 1967).

Our ACP coating, modified from established protocols based on alkaline-treated titanium catalysis (Kim et al., 2013), forms a resilient solid phase in otherwise non-adhesive cell culture plastics. Importantly, using serum-containing media stabilizes the coating, while using chemical-defined media results in more calcium release into the liquid phase of the culture. For this reason, we recommend caution in using this system in non-serum-containing experiments, as the calcium levels could be toxic for non-prostate epithelial cell lines.

All prostate epithelial-derived cells used in this study showed a high binding affinity to the ACP-coated surfaces, and enhanced proliferation, even when cultured in media supplemented with charcoal-dextran stripped serum, as a model for androgen depletion. These results are particularly important since androgen deprivation therapy (ADT) remains the standard for localized prostate cancer therapeutic care. As the tumor recedes after the initial interventions, residual disease can proliferate again, eliciting sequential patient exposure to ADT (El Badri et al., 2019). Unfortunately, some patients develop tolerance to treatment and evolve into a castration-resistant phenotype (reviewed by Hatano and Nonomura, 2023; and Kumar et al., 2022). Our results carry the dangerous implication that the presence of bioavailable calcium alone can elicit androgen independence, which could mediate, in part, the rise of castration resistance. Since ADT is a systemic treatment, it affects the bone as well. Cancer treatment-induced bone loss is a well-characterized result of hormonal ablation, with a substantial increase in bone fractures in patients on ADT (El Badri et al., 2019; Mohamad et al., 2017). Further, factors related to worsening bone health are the primary indicators of poor prognosis related to bone metastasis (Bargiota et al., 2020; Chi et al., 2016; Fizazi et al., 2015).

In agreement with our enhanced proliferation observations, ACP-coated culture elicits transcriptomic induction related to cell cycle progression, oxidative phosphorylation, and transcriptional elongation in both non-tumorigenic and early-adenocarcinoma prostatic epithelial cell lines (RWPE and LNCaP, respectively). Interestingly, our recent study on genome architecture changes in prostate cancer progression (San Martin et al., 2022) previously characterized that some of these genes are located in a B-type compartment in normal epithelium, as defined by chromosome conformation capture (Hi-C).

Since B compartments correlate with a heterochromatin identity, transcriptional induction of these genes might imply a change in chromatin architecture, similar to the one observed in the transition to adenocarcinoma or metastatic cells. Further evaluation of these genes against the Cancer Genome Atlas dataset for prostate cancer (PRAD) revealed that CDK1 and DEPDC1B expression is concomitant, correlating with a higher incidence of biochemical recurrence and increased Gleason score, which in turn signals a potentially poor prognosis.

Recent understanding of the importance of bone health in preventing metastatic progression and improving patient outcomes has given rise to combination therapies that foster bone homeostasis (Heidenreich et al., 2014; Roviello et al., 2022; Tsaur et al., 2018). However, molecular and genomic changes during prostate cancer progression and the subsequent rise of treatment resistance should influence therapy choice. (Davies et al., 2019). The ACP-coating system developed in this study is cost-effective and scalable. It is a simple way to incorporate bioavailable calcium in studies that require modeling the osteogenic bone niche. Finally, our results revealed that the presence of amorphous calcium is enough to rescue prostate epithelial cells from androgen removal *in-vitro*. Together, these findings highlight the importance of incorporating biologically relevant models that mimic the reactive bone microenvironment accompanying prostate cancer metastasis.

## Materials and Methods

### Titanium plates

Titanium Foil, 0.89mm thick, 99.7% metal basis (Alfa Aesar, AA10399GW) was cut into 2 × 2 cm plates, using an OMAX2626 JetMachining Center. A small triangular tab was included in the upper left corner of the design to facilitate identification and orientation of of the activated/exposed surface during coating.

### ACP coating

#### Titanium plates

Coating of titanium surfaces was performed according to (Kim et al., 2013) with modifications. Briefly, activation of the titanium surface was achieved by incubating four titanium plates (16 cm^2^ total surface area) in 500 ml of potassium hydroxide, at 37°C, inside a tightly closed, one-liter, polypropylene bottle (Nalgene, 75007300), for 24 hours.

After surface activation, bearing in mind to keep hydroxide-exposed surfaces up, plates were washed in three changes of miliQ water, 20 ml each, for 5 minutes under orbital shaking. Plates where then transferred, in pairs, to clean, one-liter, polypropylene bottles for ACP deposition.

In turn, ACP solution was prepared in one-liter batches, adding the following reagents stepwise: sodium chloride, potassium chloride, magnesium chloride (hexahydrated), calcium chloride (dihydrate), sodium bicarbonate and sodium phosphate (dibasic, hydrate). The sudden addition of the reagents, or an alteration of the order, caused premature precipitation. Thus, each salt was allowed to dissolve fully before adding the next one. (Table S4 as per Kim et.al (Kim et al., 2013)). Nine hundred milliliters of ACP solution were added to the polypropylene bottles, containing two titanium plates, activated side up (total surface area 8 cm^2^). Bottles were tightly closed and incubated in a 37°C oven for 24 hours. After the 24-hour incubation, two thirds of the ACP solution was collected from the incubation chambers and replaced with freshly made solution. This process was repeated two more times, for a total of 4 days incubation. All ACP solution was pooled together (Activated ACP), for usage in the coating of glass and plastic cell culture vessels.

After the four-day incubation, titanium plates were removed from the chamber, washed with an excess of milliQ water, for 30 minutes, under orbital shaking, and allowed to air dry for at least one day before sterilization (Figure 1A). Coating was verified as a white, powder deposit, and via SEM of each batch.

#### Bacterial plates

Activated ACP solution was added to plastic vessels that are non-adhesive to mammalian cells such as plastic petri dishes (Fisherbrand FB0875713. 15 ml per plate), or non-treated 6 well plates (VWR 10861-554. 5 ml/well) and incubated for four days in a 37°C oven. After incubation, ACP solution was aseptically removed and collected for filtering, while plates were washed with an equivalent volume of milliQ water, with orbital shaking, for 30 minutes. Coating was visually verified as a white powder deposit.

#### Glass coverslips

Square glass coverslips were flame-sterilized inside the biosafety cabinet, and installed in six well-plates, one coverslip per well. Coverslips were washed twice with 5 ml of PBS, and allowed to thermally stabilize inside a 37°C, 5% CO2 incubator for at least an hour prior to usage. For ACP coating, PBS was removed, and substituted with 5 ml of activated ACP, per well. Plates where then incubated in a 37°C oven for four days. After ACP incubation, plates were washed with three, 10-minute incubations in sterile milliQ water (5 ml each).

All ACP-coated vessels were sterilized under ultraviolet light, inside a biosafety cabinet, for 2 hours and stored in sterile polypropylene zip lock bags until use.

#### Activated ACP disposal

Activated ACP precipitates readily and is pervasive on most plastic and glass surfaces. The high nucleation efficiency of ACP could have a deleterious effect in the water supply and increase mineral deposition in water lines. For this reason, activated ACP was filtered through 0.2-micron membranes (Corning 430769), which significantly reduces the amount of salt aggregates in the solution, before disposal.

### X-ray diffraction (XRD) analysis

Deposited ACP was collected from 20, ten-centimeter, petri dishes. Briefly, 100 microliters of chemically pure water were added to a plate and the ACP was carefully scraped from the surface using a lifter blade scrapper (Celltreat, 229316). The 100 microliters were collected and used in subsequent plates until saturation was reached. About 500 microliters of ACP suspension were collected in total. Powdered ACP was collected by centrifugation at 5000 rpm for 10 min at room temperature. The powder was allowed to air dry overnight before X-ray diffraction analysis.

X-ray diffractograms were generated using random powder mounts and a Brüker Model D8 with Ni-filtered, Cu Kα radiation. The x-ray diffractometer operating parameters were set to 40 kV and 40 mA with a scan range of 10 to 50°2θ, and a count rate of 2 s per step. XRD analysis was performed on three ACP-coated glass coverslips and on powdered ACP, scrapped from 20 coated petri dishes. Glass coverslips were analyzed by placing them directly into the XRD instrument. The powder was suspended in water and the suspension deposited on a glass slide (creating a powder mount). After drying, the slide was placed into the XRD instrument and analyzed. Powder mounts of the mineral apatite and reagent grade NaCl (halite) were also analyzed as controls.

### Cell Culture

Prostate epithelial cell lines RWPE (Normal epithelium), LNCaP (early adenocarcinoma) and VCaP (bone metastatic) were obtained from ATCC (Cat. No CRL-11609, CRL-1740 and CRL-2876, respectively), through the Physical Sciences Oncology Network Bioresource Core Facility. All cells were cultured in T75 filtered flasks (Corning 430641U). RWPE was cultured in keratinocyte SFM, supplemented with EGF and bovine pituitary extract (Gibco 17005042), and sub cultured by incubation for 2 minutes in 1 ml of 0.25% Trypsin-0.53mM EDTA (Gibco, 25200072) at 37°C, followed by neutralization with defined trypsin inhibitor (Gibco R007100) to a ratio of 1:20 enzyme to inhibitor. LNCaP media was composed of RPMI (Gibco, 11875119) supplemented with 10% FBS (Corning 35-010-CV), glucose (Corning, 25037CI), HEPES (Gibco, 15630080) and sodium pyruvate (Gibco, 11360070) to match the concentrations suggested by ATCC (4.5 g/L glucose, 2.383 g/L HEPES and 0.11 g/L sodium pyruvate). Media used for VCaP culture was 10% FBS, in DMEM: F12 Ham 1:1 basal media.

MDAPCa2a cells were a kind gift from Dr. Nora Navone (MDAnderson Cancer Center, Houston TX). Cells were cultured in the same type of T75 vessel, coated with FCN coating mix (Athena Biologicals Cat. No. 0407), according to the manufacturers protocol. Media for MDAPCa2a cells was composed of BRFF-HPC-1 base media (Athena Biologicals Cat.No. 0403), supplemented with 20% FBS.

All cell culture media was supplemented with 1X penicillin/streptomycin (Gibco 15140122). Media formulae as described were used for culture in ACP-coated surfaces.

For trypsinization from ACP-coated petri dishes (10 cm), media was aseptically removed, and the cell monolayer was briefly rinsed with 5 ml of 5mM EDTA solution in HBSS. Cells were then exposed to a brief trypsin wash using 0.25% trypsin - 0.53mM EDTA (Gibco), supplemented to a final EDTA concentration of 1.3 mM. Cells were further incubated in another aliquot of EDTA-supplemented trypsin for 2 min in a 37°C, 5% CO_2_ incubator. After incubation, with trypsin still on the plate, plates were vortexed for 15-20 seconds, using a Vortex Genie equipped with a flat surface adaptor (setting No.3).

Enzymatic dissociation was then neutralized by adding 7 ml of full media, followed by centrifugation (5 min, 800xg). Viability and cell number was evaluated via trypan blue exclusion using a CytoSMART cell counter. This protocol was used for both subculture, and RNA-protein sample collection.

#### In-vitro androgen treatment

For supplementation of androgen in our cell culture experiments, a batch of BRFF-HPC-1 base media (Athena Biologicals) was purchased, custom-made to a final concentration of 0.5 μg/ml of dihydrotestosterone and used as stock solution. Complete media for LNCaP, VCaP and MDAPCa2a was prepared as previously described, substituting FBS with an equivalent volume of HyClone charcoal/dextran treated FBS (Cytiva/HyClone Cat.No. SH30068.03). 2.9 ml of modified BRFF-HPC-1 media was added per 50 ml of total media for a final DHT concentration of 100 mM. An equivalent volume of BMZERO media (Athena Biologicals Cat. No. 0401) was used to supplement negative control media.

Given that the RWPE cell line grows in serum-free media, for these cell lines the only supplementation necessary was modified BRFF-HPC-1 media or BMZERO control, as previously described.

#### Cell culture on titanium plates: sample preparation for electron microscopy studies

Five hundred thousand cells (VCaP, LNCaP) were seeded per sterile ACP-coated titanium plates, housed in non-treated 6 well plates, using 5 ml of cell-specific media. Cells were cultured for 4 days, with a change of media at 48 hours. Cells were fixed *in-situ* by removing 2 ml of media and adding 3 ml of 8% paraformaldehyde (Alpha Aesar, 473479M), for a final concentration of 4%, and incubated for 20 min. at room temperature. After fixation, the media/fixative was removed and plates were washed three times in 5 ml of sterile PBS, under orbital shaking. Plates were then dehydrated in a series of graded ethanol (three washes each of 70%, 95%, 100%), and allowed to air dry.

### RNA Purification and RNA-seq

RNA was purified from five million-cell pellets, using the Rneasy Plus kit (Qiagen, 74034), according to the manufacturer’s instructions, using the RLT lysis buffer supplementation with β-mercapthoethanol, as suggested. Samples were homogenized using QiaShredder columns (Qiagen, 79656), before genomic DNA removal and purification steps. RNA quality was assessed using a NanoDrop spectrophotometer.

RNA-seq was performed by Genewiz (South Plainfield, NJ) using PolyA selection for mRNA species.

### RNA-seq data processing. Differential Expression analysis and Gene ontology

RNA-seq data was processed according to previously reported pipelines. Briefly, fastq files were processed with BBDuk tool (https://github.com/kbaseapps/BBTools), trimming adapters (ktrim=r k=23 mink=11 hdist=1), and low quality reads (lower than 28: “qtrim=r trimq=28”). The reads were then aligned to the reference genome hg19 using STAR aligner (https://github.com/alexdobin/STAR) with (‘--outFilterScoreMinOverLread’ and ‘ –outFilterMatchNminOverLread’ parameters set to 0.2). Mapped reads were arranged via their genomic coordinates and feature counts generated with HTSeq-Counts (https://github.com/simon-anders/htseq).

Differential gene expression analysis between experimental conditions was evaluated using DESeq2 (https://bioconductor.org/packages/release/bioc/html/DESeq2.html) and genes defined as up/down regulated for each of the comparisons, were considered for gene ontology analysis based on a log2 fold change higher/lower than a factor of 1. Gene lists were analyzed using Metascape (Zhou et al., 2019). Full gene lists and enrichment results are available in Data S1. Gene ontology results were analyzed for commonalities among comparisons. Finally, batch adjusted counts for genes of interest were log2 normalized with a pseudo count of 1. Gene expression was plotted as a heatmap per sample/condition using the Seaborn python package (https://seaborn.pydata.org/).

### Protein collection and Western Blot

Pellets containing five million cells were collected by centrifugation at 1000 g for five minutes. After removing the supernatant, pellets were lysed in 300 μl of RIPA buffer (Thermo Scientific 89900) containing protease/phosphatase inhibitors and EDTA (GenDepot, P3100001 and P3200001), incubating on ice, for 10 minutes. 0.75 U of micrococcal nuclease (Thermo Scientific ENO0181) were added to each sample, and calcium chloride was added to a final concentration of 1 mM. Samples were then then incubated for 15 min at 37 °C, 15 min at 68°C, and on ice for 10 min, and stored at -20°C for a maximum of two weeks.

For Western blot, samples were thawed on ice and centrifuged at 14,000xg for five minutes at room temperature. The supernatant was transferred to a clean tube and the pellet was resuspended in 40 μl of 1X NuPage LDS sample buffer (Invitrogen NP0007) supplemented with DTT according to the manufacturers’ protocol. Two hundred microliters of supernatant were concentrated by acetone precipitation, adding 800 μl of ice-cold acetone and incubating at -20°C for two hours. Protein was collected by centrifugation at 14,000xg for 10 min at room temperature. The supernatant was discarded, and the pellet resuspended in 1X NuPage LDS buffer before electrophoresis.

All protein samples (membrane pellets, raw supernatant and acetone precipitated), were denatured at 70°C for 10 minutes, allowed to cool to room temperature and resolved using Bolt MES-SDS running buffer on NuPAGE 4-12% Bolt Bis-Tris mini protein gels (Invitrogen, NW04127BOX) and further transferred to methanol-activated PVDF membranes (MilliporeSigma™ Immobilon™-FL PVDF, IPFL00010) using an Invitrogen Bolt Mini Module and Bolt Transfer Buffer (Invitrogen, BT0006). Transfer was conducted at room temperature, for one hour, at 20 V.

Membranes were blocked with Intercept-TBS Blocking Buffer (LiCor 927-66003), for one hour at room temperature and exposed to 15 ml of a primary antibody solution (Rabbit polyclonal to GAPDH, Abcam ab9485), prepared at a 1:2500 ratio (15 ml) on Intercept-TBS antibody diluent (LiCor 329-66003), overnight at 4 °C, with horizontal shaking. After three washes with TBST, membranes were then exposed to IRDye 680CW Goat anti Rabbit antibody (LiCor, 92668071, 1:20,000), for 1hr, prepared in the LiCor diluent, supplemented with 0.01% SDS, in the dark with gentle shaking. After three washes with TBST (in the dark), membranes were imaged using an Odyssey CRx infrared scanner (LiCor).

We found that protein was not present in the acetone precipitated supernatant (Figure S1F) but that protein detection instead required solubilizing the membranous pellet, where the protein had aggregated due to the presence of excess calcium (Figure S1G).

### Scanning Electron Microscopy

ACP-coated titanium plates were imaged in the using a Hitachi TM3030 scanning electron microscope at an accelerating voltage of 15kV, a working distance between 6800 and 7000 microns, at a magnification of x5000.

### Evaluation of Calcium Leaching

Two ACP-coated petri dishes from at least three different ACP coating batches were incubated with 15 ml of full RWPE, LNCaP and VCaP media for four days, at 37°C, 5% CO_2_. After incubation, media was aseptically collected, pooled, filtered through 0.2-micron PVDF filters (low protein binding) and stored at -20°C. Samples were diluted 1:10 with chemically pure water before analysis.

Calcium leaching was measured at the University of Tennessee Water Quality Core Facility using ion chromatography (IC) using a Thermo-Scientific®/Dionex® ICS-1100 (cations) system with background suppression for low detection limits.

## Acknowledgments

The authors would like to acknowledge Jaydeep Kolape and the University of Tennessee Advanced Microscopy and Imaging Center for instrument use, and scientific and technical assistance with SEM.

## Data Availability

All gene expression data for prostate cancer cell lines grown with and without testosterone on control and ACP surfaces are available at GEO accession number GSE227511 https://www.ncbi.nlm.nih.gov/geo/query/acc.cgi?acc=GSE227511.

